# Neural variability reliably and selectively encodes pain discriminability

**DOI:** 10.1101/2025.02.26.640289

**Authors:** Libo Zhang, Xinyi Geng, Li Hu

## Abstract

Neural activity varies dramatically across time. While such variability has been associated with cognition, its relationship with pain remains largely unexplored. Here, we systematically investigated the relationship between neural variability and pain, particularly pain discriminability, in five large electroencephalography (EEG) datasets (total N = 489), collected from healthy individuals (Datasets 1–4) and patients with postherpetic neuralgia (PHN; Dataset 5) who had received painful or nonpainful sensory stimuli. We found robust correlations between neural variability and interindividual pain discriminability. These correlations were (1) replicable in multiple datasets, (2) pain selective, as no significant correlations were observed in nonpain modalities, and (3) clinically relevant, as they were partly disrupted in patients with PHN. Importantly, variability and amplitude of EEG signals were mutually independent and had distinct temporal and oscillatory profiles in encoding pain discriminability. These findings demonstrate that neural variability is a replicable and selective indicator of pain discriminability above and beyond amplitude, thereby enhancing the understanding of neural encoding of pain discriminability and underscoring the value of neural variability in pain studies.

## Introduction

Cognitive neuroscience studies typically characterize neural activity in terms of its amplitude. For example, the most common electroencephalography (EEG) metrics in pain studies involve the amplitude of nociceptive evoked potentials, such as the N2 and P2 components [1–5]. Although this approach has remarkably advanced the understanding of pain processing, it overlooks another critical aspect of neural activity: its variability. Importantly, in some fields, such as perception and memory, neural variability has been increasingly recognized as having a functional role [6–8]. A positive correlation has been found between increased variability in functional magnetic resonance imaging signals within the human motion complex and enhanced discrimination of visual motion direction [9]. In addition, neural variability in EEG signals has also been associated with interindividual differences in reaction time during face-recognition tasks [10]. Thus, neural variability may provide new insights into how the brain represents pain information beyond the traditional amplitude-based approach.

Previous amplitude-based research has greatly contributed to revealing the neural encoding of pain [11–14]. Pain variations at intraindividual and interindividual levels have been associated with the amplitudes of N2, P2, and gamma-band oscillations [3,15–17]. However, many previous studies faced a critical issue: a lack of pain selectivity. EEG responses often correlate with perceptual variation in nonpain sensory modalities as well [15,18]. Interestingly, pain discriminability, the ability to distinguish between two or more painful stimuli, appears to be selectively encoded by the amplitude of event-related potentials (ERPs) [19,20]. This selective encoding is particularly noteworthy. For one thing, ERP components such as N2 and P2 have a much higher signal-to-noise ratio than other responses, such as gamma-band oscillations [21,22]. For another, pain discriminability is clinically relevant. Impaired pain discriminability has been observed in patients with chronic pain [23,24], and higher pain discriminability can predict a better effect of pain treatment [25,26]. In reality, however, pain discriminability remains severely under-investigated [15,16,20,27,28].

In contrast to the amplitude-based approach, few, if any, studies have systematically examined how neural variability contributes to the neural encoding of pain, particularly pain discriminability. Many fundamental questions remain unanswered. First, it is unclear whether neural variability reflects pain. Currently, only a handful of studies have *indirectly* associated neural variability with some aspects of pain, such as with temporal summation of pain [29,30]. The relationship between neural variability and pain, particularly pain discriminability, thus remains largely unexplored. Second, it remains to be tested whether the association between neural variability and pain discriminability is replicable. The replication crisis has called into question even classic findings in biological sciences [31–34]. The replicability of the association must be proven before the role of neural variability can be asserted with confidence. Third, it is unclear whether and how the role of neural variability in encoding pain discriminability differs from that of the amplitude of neural activity. Given the association between ERP amplitude and pain discriminability [20], this question is key to establishing the independent role of neural variability as an indicator of pain discriminability, above and beyond the amplitude of neural activity. Fourth, it has yet to be determined whether neural variability is selectively related to pain discriminability. A mechanistically insightful and clinically useful neural indicator should be pain selective [12]. Otherwise, the indicator could be driven by stimulus-general confounders and provide limited insights into the neural mechanisms.

To address these questions, we systematically examined the relationship between neural variability and pain, with a particular focus on pain discriminability, using five large EEG datasets (total N = 489; see Figure 1). Datasets 1–4 were collected from healthy participants exposed to painful laser stimuli and nonpainful tactile, auditory, and visual stimuli of varying intensities, while Dataset 5 was collected from patients with postherpetic neuralgia (PHN; a type of chronic pain) receiving painful laser stimuli. Since ERP amplitude has been associated with pain discriminability [20], we specifically focused on evaluating the independence of neural variability from ERP amplitude in encoding pain discriminability. Although our focus was on pain discriminability, we also explored the relationship between neural variability and intraindividual pain perception and interindividual pain sensitivity.

**Figure 1.**
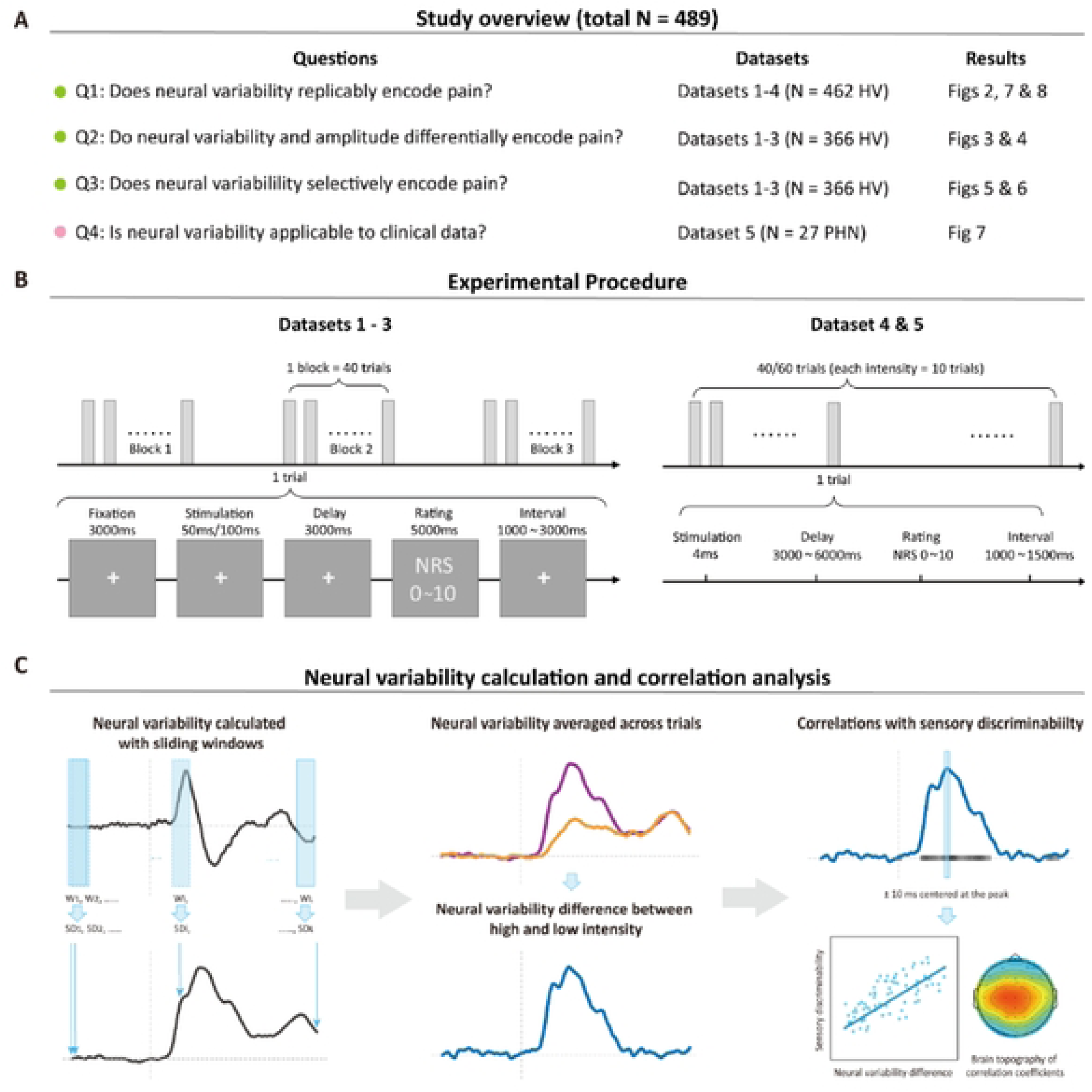
Study overview, experimental procedures, and neural variability. **(A)** Overview of the study. Participants in Datasets 1-4 were healthy volunteers (HV), whereas participants in Dataset *5* were chronic pain patients with postherpetic neuralgia (PHN). **(B)** Experimental procedures. All participants in Datasets 1−3 received a total of 120 stimuli across four sensory modalities (pain, touch, audition, and vision) presented in a pseudorandom order in three blocks. After each stimulus, participants rated the perceived intensity using a numerical rating scale (NRS), which ranged from 0 (no sensation) to 10 (the strongest imaginable sensation). Participants in Datasets 4 and *5* received painful laser stimuli at multiple levels of intensity in a pseudorandom order. For each intensity, ten pulses were delivered. After each stimulus, participants rated the perceived intensity using the NRS. (C) Calculation of temporal SD as a measure of neural variability. Temporal SD was derived from windowed EEG responses evoked by each stimulus. A series of 100-ms sliding windows with a 1-ms step was employed. SD difference (ΔSD) was then calculated by subtracting the trial­ averaged SD in the low-intensity condition from that in the high-intensity condition. Correlation analyses were conducted to evaluate the relationship between neural variability and sensory discriminability.

## Results

### Neural variability reliably encodes pain discriminability independent of amplitude

We employed temporal standard deviation (SD) calculated with 100-ms sliding windows as the primary metric for neural variability in this study (Figure 1C) and verified the robustness of our findings with windows of other lengths. In Dataset 1, pain stimuli with higher intensity evoked greater variability in EEG responses, particularly over the central region of the brain (Figure 2A). Additionally, pain ratings were higher in the high-intensity condition than in the low-intensity condition (*t*_(140)_ = 17.08, *P* < 0.0001, Cohen’s *d* = 1.44). To determine the relationship between neural variability and pain discriminability (quantified as the difference in pain ratings between high- and low-intensity conditions), we then correlated them at every time point and corrected the *P* values with false discovery rate (FDR) correction. Significant correlations were identified (i.e., time points with gray-shaded bars) across participants between neural variability differences (i.e., temporal SD difference, ΔSD) in the high- and low-intensity conditions and pain discriminability in Datasets 1 (Figure 2B). Scatter plots further indicated no gross outliers or nonlinearity affecting the corrections around the peak (±10 ms centered at the peak) of ΔSD (Dataset 1: *r* = 0.48, *P* = 1.54×10^-9^; Figure 2D).

**Figure 2.**
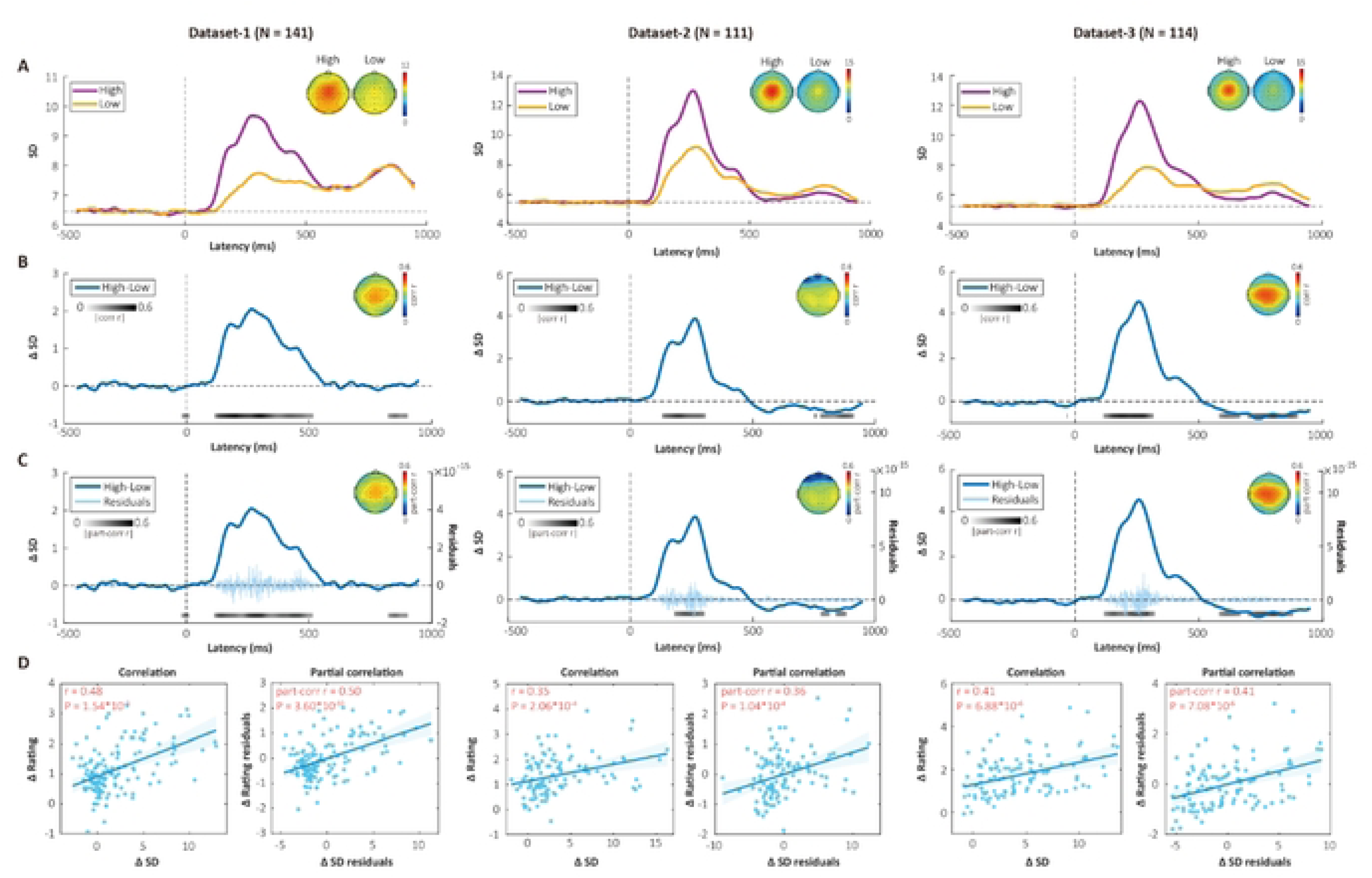
Neural variability reliably correlated with pain discriminability. **(A)** Neural variability assessed by a variance-based measure, temporal SD of electroencephalography (EEG) responses evoked by high-intensity (violet) and low-intensity (gold) stimuli in Datasets 1−3. **(B)** Point-by-point correlations between neural variability differences (high-low, ΔSD) and rating differences (high-low, ΔRating)in the three datasets. **(C)** Point-by-point partial correlations between ΔSD and ΔRating while controlling for amplitude differences (high-low, ΔAmpli1ude). The light-colored “bursty” curve represents the subject-averaged residuals of ΔSD after regressing out ΔAmplitude at each time point. **(D)** Correlations and partial correlations between values around the peak of neural variability(± 10 ms) and ΔRating. Partial correlations were calculated while controlling for ΔAmplitude. Note that the gray bars represent Pearson’s *r* values at time points where significant correlations were observed after false discovery rate correction. Part.corr = Partial correlation. Error bars in **D** are 95% confidence intervals.

Statistically speaking, SD is inherently correlated with mean values, which was verified in Dataset 1 (Supplementary Figure 1B). A previous study has shown that mean ERP amplitude differences between high- and low-intensity conditions also correlate with pain discriminability [20], which was also confirmed here by applying a 100-ms sliding window analysis to the ERP waveforms (Supplementary Figure 1C&D). ERP amplitude thus could potentially confound the relationship between neural variability and pain discriminability described above. To rule out this possibility, we conducted three sets of control analyses. First, we performed partial correlations between ΔSD and pain discriminability, controlling for mean amplitude differences (ΔAmplitude). Significant partial correlations between ΔSD and pain discriminability were observed (Figure 2C&D; partial *r* around the peak = 0.50, *P* = 3.60×10^-10^). Notably, the time points showing significant partial correlations nearly perfectly overlapped with those showing simple correlations. Interestingly, amplitude correlated with pain discriminability after accounting for neural variability as well (Supplementary Figure 1E&F), suggesting that neural variability and amplitude were mutually independent in their roles in encoding pain discriminability. Second, we repeated the foregoing correlation analysis with an information-theory-based measure of neural variability: permutation entropy (PE). PE quantifies the complexity of a time series by evaluating the ordinal relations between values in the time series rather than their absolute numerical values [35], and is thus less affected by amplitude. Significant correlations were again observed between PE differences (ΔPE) and pain discriminability (Supplementary Figure 2). As a third control analysis, we subtracted trial-averaged ERPs from each trial before calculating temporal SD, that is, analyzing induced responses instead of evoked responses [36]. Still, significant correlations were observed between neural variability differences (assessed by temporal SD and PE) of the induced responses and pain discriminability in Dataset 1 (Supplementary Figure 3). Altogether, these findings lend strong support to the statement that neural variability encodes pain discriminability independently of amplitude.

To test the robustness of our findings, we conducted further sensitivity analyses, examining the influence of metrics for pain discriminability and varying window sizes for computing neural variability. Instead of simple differences in pain ratings, we quantified pain discriminability with the area under the curve (AUC) derived from signal detection theory (for a detailed explanation of AUC, please refer to [20]). Aligning with our previous observations, neural variability exhibited a noteworthy correlation with pain discriminability measured by AUC (Supplementary Figure 4). We then calculated neural variability using alternative time-window sizes of 50 ms and 200 ms. Our results remained consistent: pain discriminability correlated with neural variability regardless of the window sizes (see Supplementary Figure 5). These findings provide evidence for the robustness of neural variability as an indicator of pain discriminability.

To assess the replicability of our findings, we then examined the relationship between neural variability and pain discriminability in two large independent datasets. Positive correlations were observed in Datasets 2 and 3 (Figure 2B). Additionally, ΔSD around the peak exhibited significant correlations with pain discriminability in both datasets (Dataset 2: *r* = 0.35, *P* = 2.06×10^-4^; Dataset 3: *r* = 0.41, *P* = 6.88×10^-6^; Figure 2D). Further analyses once again confirmed that ERP amplitude did not explain the correlations between neural variability and pain discriminability in Datasets 2 and 3 (see partial correlation results in Figure 2C&D). Collectively, these findings underscore the replicability and reliability of neural variability as an independent indicator of pain discriminability.

### Neural variability encodes pain discriminability earlier and more efficiently than amplitude with distinct oscillatory profiles

Given that both neural variability and ERP amplitude have the independent capability to encode pain discriminability, a key question emerges: how do they differ? To address this question, we first conducted dominance analysis to separate the relative contributions of neural variability and ERP amplitude to pain discriminability. Dominance analysis is a method that decomposes the total determination coefficient (*R*^2^) of a multiple regression model into the unique *R*^2^ (or “total dominance” in dominance analysis’s terminology) contributed by each predictor [37,38]. We performed dominance analysis at each time point to evaluate the total dominance of neural variability and ERP amplitude. In Dataset 1, the full model with both SD and amplitude as predictors exhibited two peaks at around 200 ms and 400 ms, roughly corresponding to the N2 and P2 components of ERPs (Figure 3A). Neural variability played a greater role in the rising phase of the first peak, while ERP amplitude predominantly contributed to the second peak. Similar patterns could also be observed in Datasets 2 and 3. These findings suggest that neural variability and ERP amplitude differ in the time course of their encoding pain discriminability. Specifically, neural variability plays a role equal to or greater than amplitude in encoding pain discriminability around 100–300 ms after stimulus onset, while ERP amplitude plays a more important role about 300–500 ms after stimulus onset.

**Figure 3.**
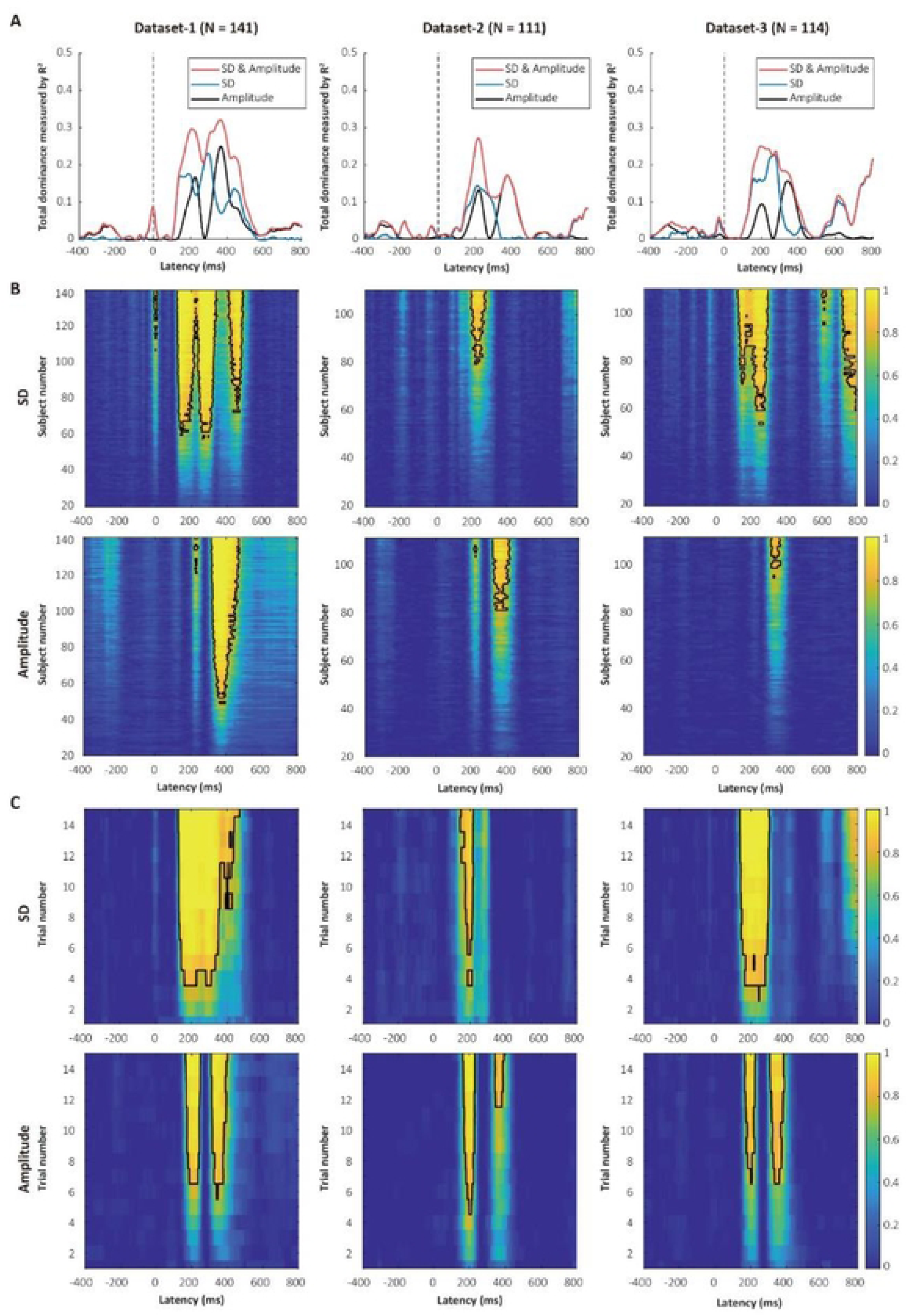
Comparisons of neural variability and ERP amplitude for encoding pain discriminability. **(A)** Relative contributions of neural variability and event-related potential (ERP) amplitude in Datasets 1−3. SD and ERP amplitude exhibited different contributions to pain discriminability over time, with the former having greater contributions around 100-300 ms after stimulus onset and the latter having greater contributions around 300- 500 ms. Curves represent total dominance measured by determination coefficient *R*^2^. **(B)** The influence of the number of subjects on the probability of detecting significant correlations between SD or ERP amplitude and pain discriminability after false discovery rate (FDR) correction in the three datasets. Color bars indicate the probability of detecting the correlations. The black contour represents the probability of significance that exceeds 80%. Using neural variability or ERP amplitude as indicators of pain discriminability may require a comparable number of subjects. These results further confirm that SD and amplitude exhibited different contributions to pain discriminability over time. **(C)** The influence of the number of trials on the probability of detecting significant correlations between SD or ERP amplitude and pain discriminability after FDR correction in the three datasets. Using neural variability as an indicator of pain discriminability requires fewer trials than using ERP amplitude.

To further characterize the differences between neural variability and ERP amplitude, we applied a resampling approach to explore whether the number of subjects influences the probability of detecting significant correlations between SD or amplitude and pain discriminability in Datasets 1–3 (see **Methods** for resampling details). We found no clear evidence that either neural variability or ERP amplitude requires a smaller number of subjects to exhibit their correlations with pain discriminability with 80% power (Figure 3B). Interestingly, we observed a difference in the time windows when comparing the required number of subjects using SD and amplitude. Specifically, with the same number of subjects, the correlations between SD and pain discriminability could be detected earlier than those between amplitude and pain discriminability. These results further confirmed that neural variability encodes pain discriminability earlier than amplitude.

Neural variability can be computed at the single trial level, but ERP amplitude is generally calculated by averaging multiple trials. We thus hypothesized that another difference between neural variability and ERP amplitude is that neural variability may require a smaller number of trials to exhibit correlations with pain discriminability. To test this hypothesis, we applied the resampling approach to evaluate the influence of number of trials on the probability of detecting significant correlations between neural variability or ERP amplitude and pain discriminability. The number of trials that could yield a significant correlation between SD and pain discriminability with 80% power was about 4 in the three datasets (Figure 3C, upper panels). In contrast, using ERP amplitude as an indicator of pain discriminability required more trials (about 7) to meet the same criteria in the three datasets (Figure 3C, lower panels). Therefore, neural variability encodes pain discriminability more efficiently than ERP amplitude.

To reveal the oscillatory profiles of neural variability and ERP amplitude, we then band-pass filtered single-trial EEG signals according to the classic frequency bands (i.e., delta, theta, alpha, and beta) and conducted a partial correlation analysis. ERP-like deflections could be found only in the delta and theta frequency bands in Dataset 1 (Figure 4). Consistent with this observation, after we controlled for neural variability, significant correlations between amplitude and pain discriminability were seen only in the delta and theta frequency bands, and not in the alpha and beta frequency bands (Figure 4). In contrast, the partial correlations between neural variability and pain discriminability after regressing out ERP amplitude were present in all four frequency bands, even in frequencies without ERP responses (Figure 4). These findings were replicated in Datasets 2&3 (Supplementary Figure 6), suggesting that neural variability and ERP amplitude had different oscillatory profiles in encoding pain discriminability.

**Figure 4.**
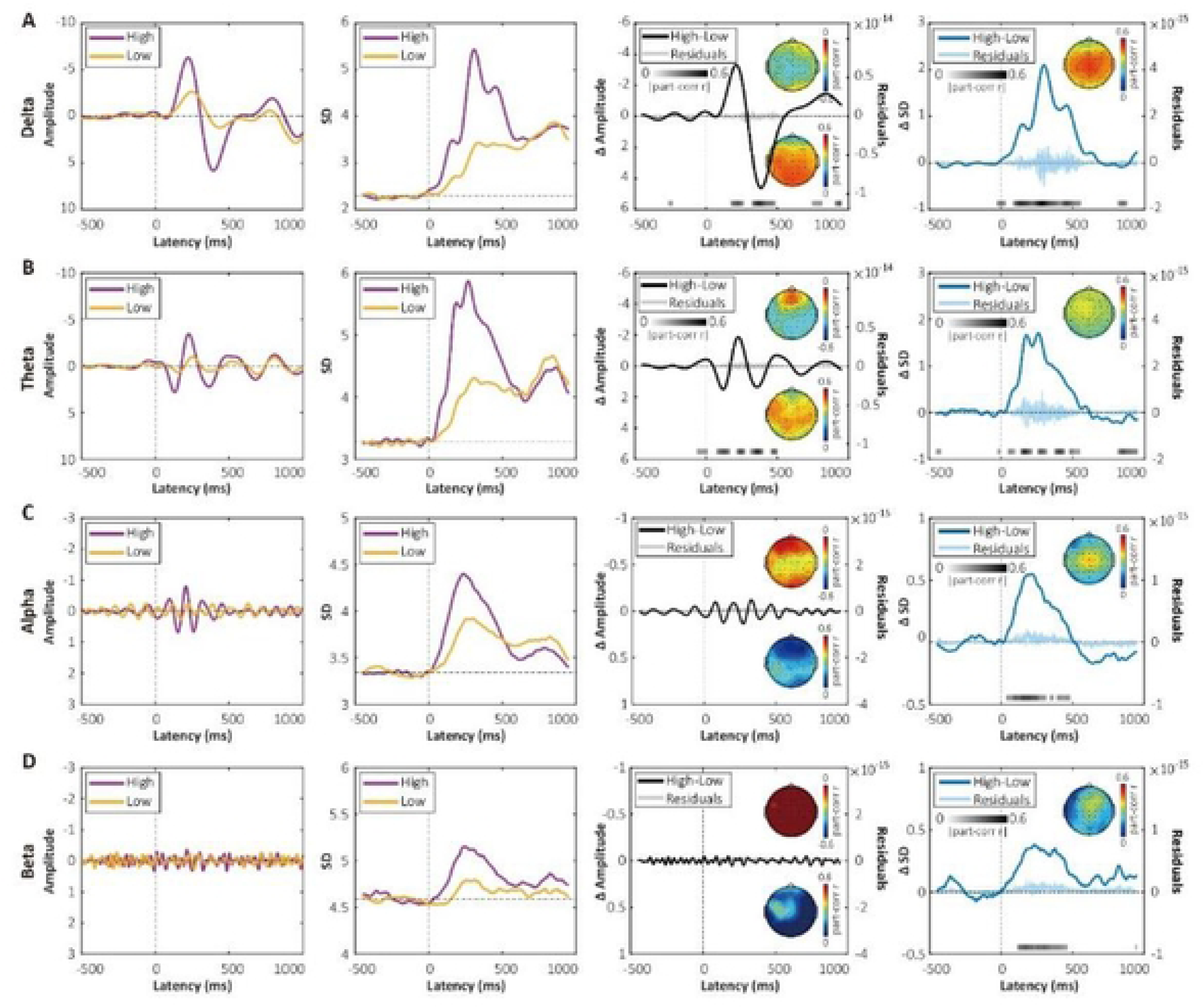
Comparisons of neural variability and ERP amplitude at different frequency bands for encoding pain discriminability in Dataset 1. **(A-D)** Event-related potential (ERP) amplitude, neural variability, and the differential values between high- and low-intensity stimuli at the delta **(A,** 1−4 Hz), theta (**B,** 4−8 Hz), alpha (**C,** 8−12 Hz), and beta **(D,** 12−30 Hz) bands, and their partial correlations while controlling for each other with pain discriminability. The two left columns show ERP amplitude and neural variability of band-pass-filtered time series averaged across trials evoked by high-intensity (violet) and low-intensity (gold) stimuli. The two right columns show point-by-point partial correlations of ERP amplitude differences (high­ low, ΔAmplitude) and SD differences (high-low, ΔSD) with pain discriminability, respectively. Note that the gray bars represent Pearson’s *r* values at time points where significant correlations were observed after false discovery rate correction. Part-corr= Partial correlation. The topographies represent partial *r* values within a± 10-ms window around the peak. For ΔAmplitude calculations in **A** and **B,** the latencies of N2 and P2 peaks were utilized, while for **C** and **D**, where clear ERPs were not observed, latencies corresponding to the minimal and maximal values within the 100-500 ms time window were used.

### Neural variability selectively encodes pain discriminability

After establishing the replicability and independence of neural variability as an indicator of pain discriminability, we proceeded to investigate its selectivity. Note that participants who provided Datasets 2 and 3 were recruited on a rolling basis and assigned to either Dataset 2 or Dataset 3 according to their pain sensitivity, which was assessed during the calibration phase. We thus combined these two datasets (denoted Datasets 2&3) in the analyses of nonpain modality data. We found no reliable evidence for correlations between neural variability and discriminability in the tactile, auditory, or visual modalities. In Dataset 1, significant but weak correlations were observed only in the tactile modality within 35–180 ms, which were before the peak of the ΔSD wave (see Figure 5A). However, no significant correlations were observed between the peak values of ΔSD and sensory discriminability after controlling for the amplitude of sensory ERPs: (1) touch: partial *r* = 0.13, *P* = 0.14; (2) audition: partial *r* = 0.14, *P* = 0.09; (3) vision: partial *r* = 0.05, *P* = 0.59 (Figure 4A–C). Moreover, in Datasets 2&3, we did not find significant correlations between neural variability and sensory discriminability in nonpain modalities: (1) touch: partial *r* = 0.06, *P* = 0.35; (2) audition: partial *r* = 0.11, *P* = 0.10; (3) vision: partial *r* = 0.09, *P* = 0.18 (Figure 5D–F).

**Figure 5.**
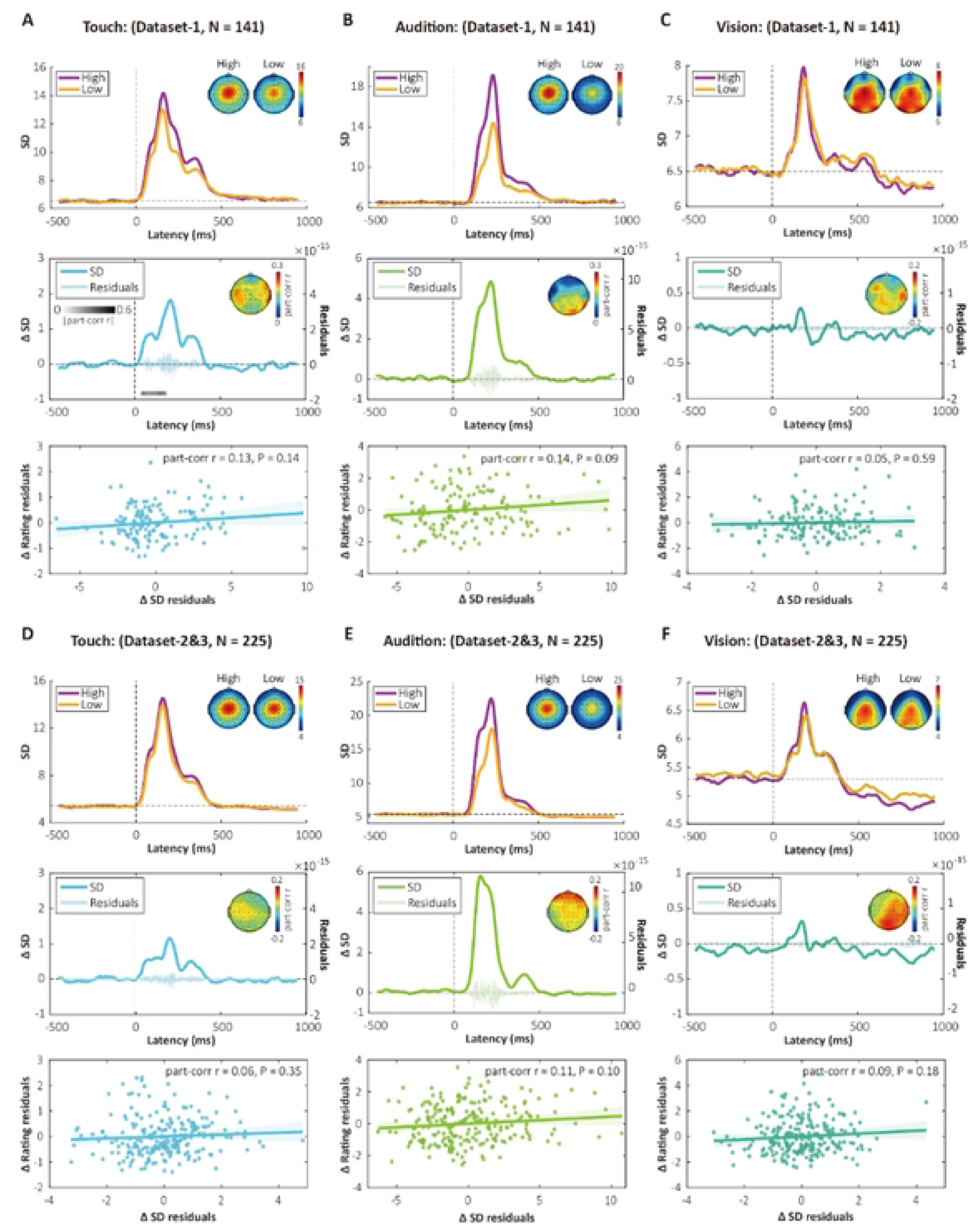
Neural variability of electroencephalography responses evoked by tactile, auditory, and visual stimuli did not reflect their respective sensory discriminability. **(A-C)** Neural variability, and its partial correlations with sensory discriminability of tactile, auditory, and visual stimuli in Dataset 1. Gray bars represent Pearson’s *r* values at time points where significant correlations were observed (false discovery rate corrected). For tactile modality, significant partial correlations while controlling for differences in event-related potential (ERP) amplitude were found at 35−180 ms, but no significance was observed around the peak of neural variability. No significance was observed for either the auditory or visual modality in partial correlations while controlling for ERP amplitude differences. **(D-F)** Neural variabilities and their partial correlations with respective sensory discriminability in Dataset 2&3. No significant partial correlations while controlling for ERP amplitude differences were observed for any of these sensory modalities. Part-corr = Partial correlation. The light-colored “bursty” curves represent the subject-averaged residuals of ΔSD after regressing out ΔAmplitude at each time point. Error bars are 95% confidence intervals.

These results offer preliminary evidence that neural variability selectively encodes pain discriminability. However, a potential confounding factor arises from the possibility that sensory discriminability, quantified as rating differences, could vary across modalities. Indeed, pain discriminability was smaller compared to auditory and visual discriminability (e.g., paired-sample *t* tests in Dataset 1: pain vs. audition: *t*_(140)_ = −11.45, *P* = 7.97×10^-22^; pain vs. vision: *t*_(140)_ = −10.31, *P* = 6.75×10^-19^; Dataset 2&3: pain vs. audition: *t*_(224)_ = - 27.90, *P* = 7.98×10^-75^; pain vs. vision: *t*_(224)_ = - 14.47, *P* = 6.12×10^-34^). A significant difference was also observed between pain and tactile discriminability in Dataset 2&3 (*t*_(224)_ = 6.08, *P* = 5.09×10^-9^). To ensure rigorous and fair comparisons, we implemented a rating-matching procedure (see **Methods** for details). This approach enabled us to identify subgroups of participants who exhibited comparable intensity ratings across pairs of sensory modalities, specifically pain versus touch, pain versus audition, and pain versus vision. Through this method, Dataset 1 yielded 70 participants for the pain–touch matching, 33 for the pain–audition matching, and 58 for the pain–vision matching. Within these matched subgroups, significant correlations between neural variability and pain discriminability were consistently observed even after controlling for ERP amplitude, while no significant correlations were found in nonpain modalities: (1) pain versus touch: pain, partial *r* = 0.49, *P* = 2.27×10^-5^; touch, partial *r* = 0.17, *P* = 0.17; (2) pain versus audition: pain, partial *r* = 0.40, *P* = 0.02; audition, partial *r* = 0.17, *P* = 0.35; (3) pain versus vision: pain, partial *r* = 0.49, *P* = 9.10×10^-5^; vision, partial *r* = −0.02, *P* = 0.87 (Figure 6A–C). Similar findings were obtained in Datasets 2&3: (1) pain versus touch: pain, partial *r* = 0.37, *P* = 1.64×10^-4^; touch, partial *r* = 0.14, *P* = 0.16; (2) pain versus audition: pain, partial *r* = 0.46, *P* = 4.05×10^-3^; audition, partial *r* = 0.01, *P* = 0.56; (3) pain versus vision: pain, partial *r* = 0.35, *P* = 0.001; vision, partial *r* = −0.03, *P* = 0.77 (Figure 6D–F). These consistent results compellingly demonstrate that neural variability selectively encodes pain discriminability.

**Figure 6.**
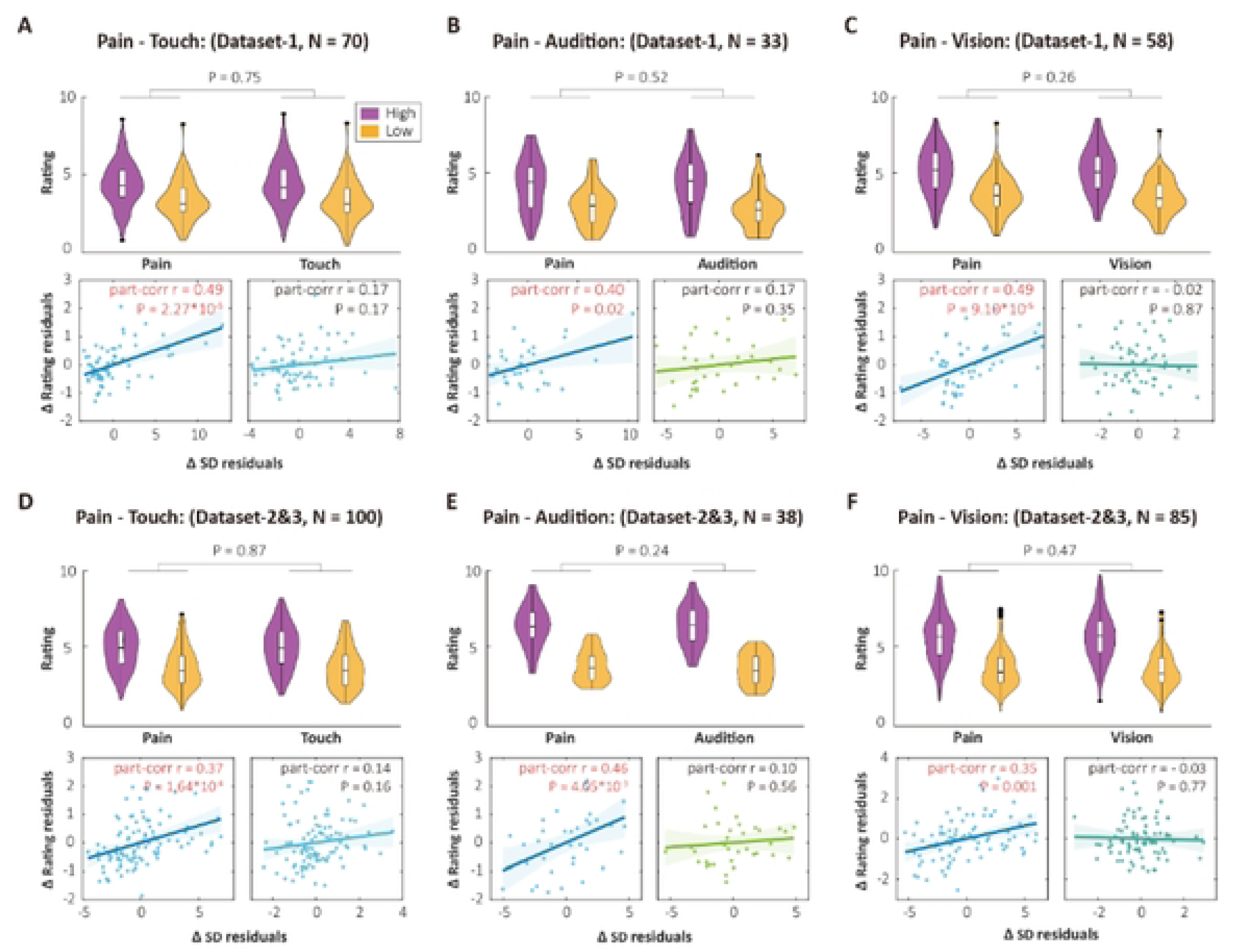
Selectivity of neural variability as an indicator of pain discriminability by matching the ratings between pain and nonpain modalities. **(A&D)** The upper panels show distributions of ratings for high- and low-intensity stimuli matched between the pain and tactile modalities in Dataset 1 and Datasets 2&3. The high-low rating differences between pain and touch were not significant after rating matching. The lower panels show partial correlations between SD differences and rating differences for the matched subset. **(B, C, E, F)** Similar figures for subsets matched between pain and auditory modalities, and between pain and visual modalities in Datasets 1 and 2&3. Significant partial correlations were found in all matched subsets for pain discriminability, but not for tactile, auditory, and visual modalities in both datasets. Part-corr= Partial correlation. Error bars are 95% confidence intervals.

### Generalizability and clinical relevance of neural variability as an indicator of pain discriminability

In Datasets 1–3, the differences in laser stimulus energy between high- and low-intensity conditions remained fixed at 0.5 J. To examine the generalizability and clinical relevance of the correlations between neural variability and pain discriminability, we conducted additional analyses on Datasets 4 and 5, in which participants received laser stimuli of multiple intensities, specifically, 2.5 J, 3.0 J, 3.5 J, and 4.0 J in Dataset 4 (N = 96, healthy subjects), and 3.0 J, 3.25 J, 3.5 J, 3.75 J, 4.0 J, and 4.25 J in Dataset 5 (N = 27, patients with PHN). Note that patients with PHN received stimulation both on a shingles-affected skin area (pain-affected) and a mirrored skin area on the contralateral side (pain-unaffected).

In Dataset 4, pain ratings were significantly different across stimulus intensities (one-way repeated-measures analysis of variance, ANOVA, *F*_(3, 380)_ = 182.47, *P* = 2.86×10^-73^; Figure 7A). Temporal SDs were significantly different within 100–500 ms in all pairs of high- and low-intensity stimuli (Figure 7A). Correlations between ΔSD and pain discriminability showed significance in four pairs of high and low stimulus intensities: (1) 4.0 J versus 3.5 J: *r* = 0.21, *P* = 0.04; (2) 3.5 J versus 3.0 J: *r* = 0.31, *P* = 0.002; (3) 3.0 J versus 2.5 J: *r* = 0.25, *P* = 0.02; (4) 3.5 J versus 2.5 J: *r* = 0.27, *P* = 0.01 (Figure 7B). Thus, the correlations between neural variability and pain discriminability may be reliably detected only when the intensity difference is moderate (e.g., 0.5 J), which may be due to a potential ceiling effect on pain discriminability when the intensity difference becomes excessively large.

**Figure 7.**
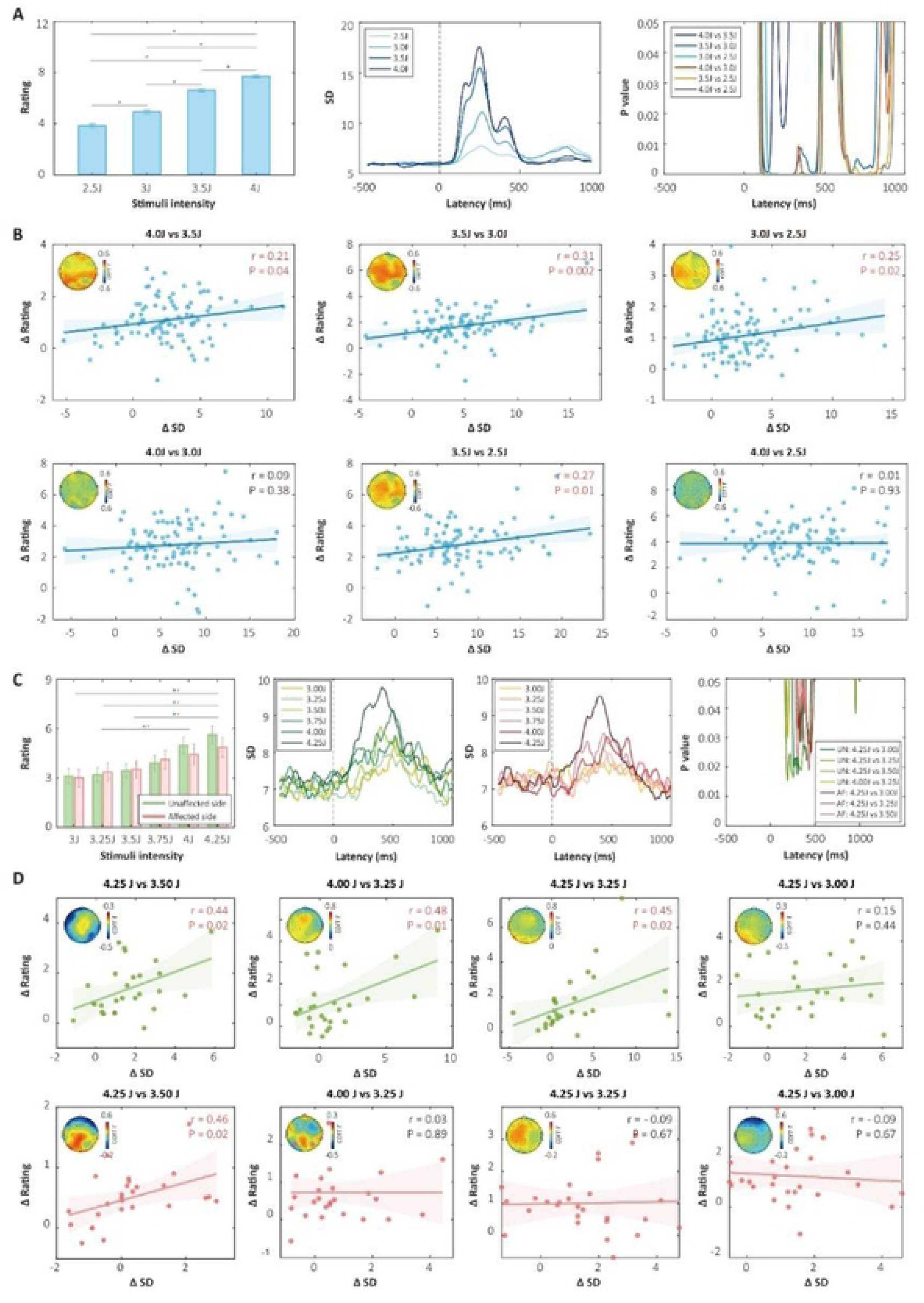
Neural variability as an indicator of pain discriminability in Dataset 4 (healthy subjects) and Dataset 5 (patients with chronic pain) with multiple levels of stimuli. **(A)** Pain intensity ratings (left panel), temporal SD (middle panel), and *P* values of paired-sample *t* tests for temporal SD between each pair of stimulus intensities (right panel) in Dataset 4 (healthy subjects, N = 96). Notably, ratings were significantly different across stimulus intensities. The right panel illustrates the time points where neural variability was significantly different between pairs of stimulus intensities. **(B)** Correlations between ΔSD around the peak and ΔRating in all pairs of stimulus intensities. Significant correlations were found in discriminating between intensities with a 0.5-J difference (upper panels), as well as between 3.5 J and 2.5 J (middle lower panel). **(C)** Pain intensity ratings (left panel), temporal SD in the pain-unaffected and pain-affected sides (middle two panels), and *P* values of paired-sample *t* tests for temporal SD between pairs of stimulus intensities (right panel) in Dataset 5 (patients with postherpetic neuralgia, N = 27). Paired-sample *t* tests revealed significant differences for most stimulus pairs with a difference exceeding 0.5 J on both sides. Significance is highlighted in green for the unaffected side and red for the affected side (left panel). Temporal SDs showed significant differences in 7 pairs of stimulus intensities (right panel). **(D)** Correlations between ΔSD and pain discriminability were assessed for both the unaffected (upper panels) and the affected (lower panels) side. Significant correlations were observed in three pairs of intensities on the unaffected side and one pair of intensities on the affected side, indicating a disruption of the correlations in the affected side. Error bars in **A** and **C** are standard errors of means, and error bars in **B** and **D** represent 95% confidence intervals.

In Dataset 5, stimulus intensity had a significant influence on ratings, while stimulation side and the interaction between stimulus intensity and side were not significant (two-way repeated-measures ANOVA, stimulus intensity: *F*_(5, 312)_ = 5.46, *P* = 7.85×10^-5^; side: *F*_(1, 312)_ = 0.28, *P* = 0.60; interaction: *F*_(5, 312)_ = 0.30, *P* = 0.92). Significant differences were found between most stimulus pairs with a difference ≥ 0.75 J on both sides (leftmost panel of Figure 7C). Significant temporal SD differences were observed in seven pairs of intensities (rightmost panel of Figure 7C). Among those intensity pairs with significant rating and SD differences, correlations between ΔSD and pain discriminability showed significance in three intensity pairs on the unaffected side (4.25 J vs. 3.5 J: *r* = 0.44, *P* = 0.02; 4.0 J vs. 3.25 J: *r* = 0.48, *P* = 0.01; 4.25 J vs. 3.25 J: *r* = 0.45, *P* = 0.02), but in only one pair on the affected side (4.25 J vs. 3.5 J: *r* = 0.46, *P* = 0.02). The associations between neural variability and pain discriminability therefore seem to have been disrupted to some extent on the chronic-pain-affected side, suggesting potential clinical relevance of neural variability as an indicator of pain discriminability.

### Neural variability reflects general intraindividual sensory perception but not interindividual sensory sensitivity

To investigate more comprehensively the relationship between neural variability and pain, we then examined whether neural variability reflects interindividual pain sensitivity and intraindividual pain perception. For interindividual sensitivity, temporal SD waves were averaged across all trials regardless of stimulus intensity and then correlated with trial-averaged ratings. After controlling for means of ERP amplitude, no significant correlations were found between trial-averaged SD and ratings for all modalities in Dataset 1 (N = 141) and Dataset 2&3 (N = 225; Figure 8A&C). For intraindividual sensory perception, correlation analyses were conducted at the single-trial level for each subject, and the Fisher *z*-transformed correlation coefficients were compared with zero using a one-sample *t* test. In contrast to the results for interindividual sensitivity, significant correlations were observed for all modalities (Figure 8B&D). These findings align with previous studies demonstrating that the amplitude of most brain responses to nociceptive stimuli, though not pain selective, reflects pain perception at the intraindividual level, while ERP amplitude per se does not encode pain sensitivity at the interindividual level [15].

**Figure 8.**
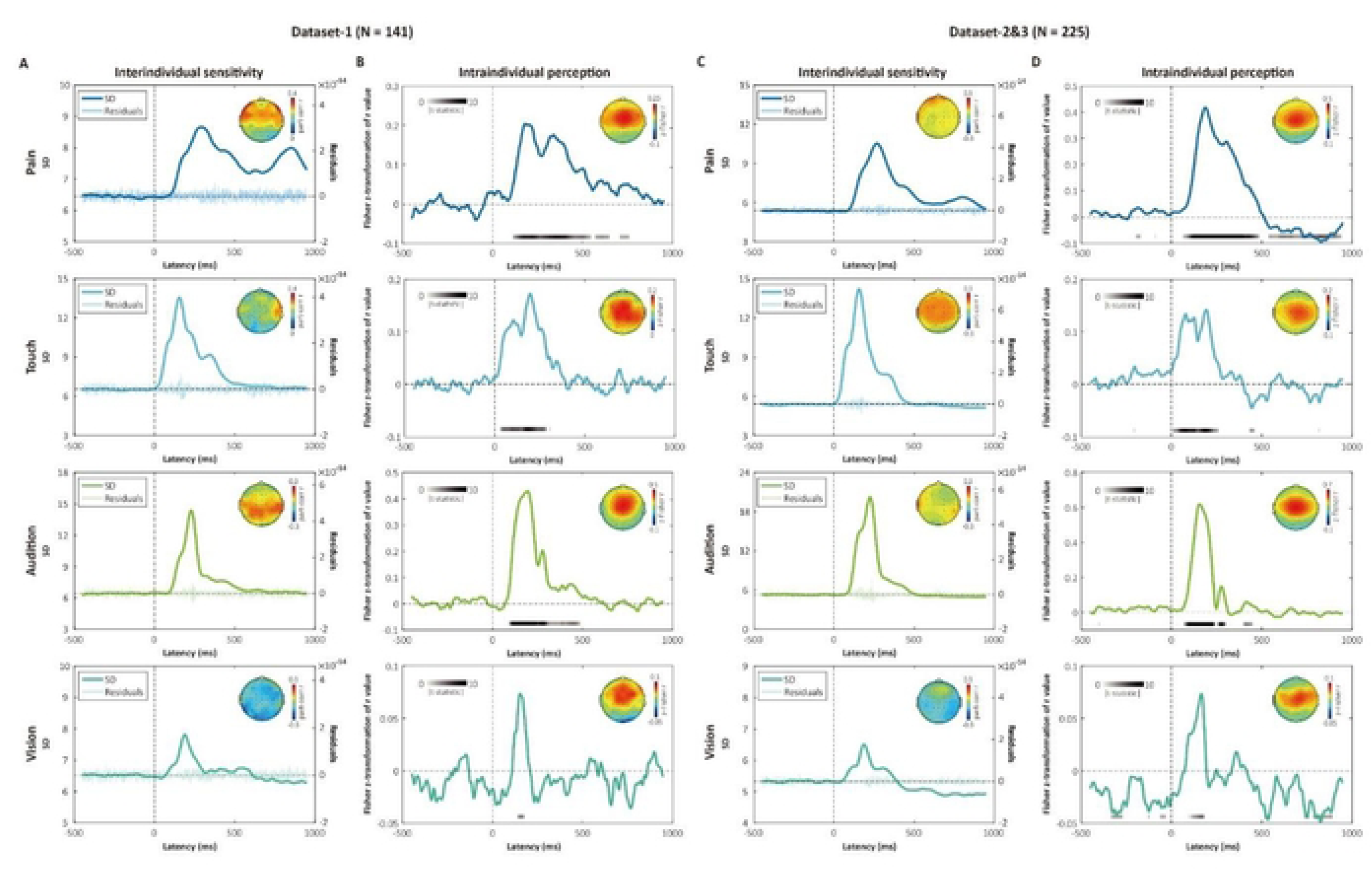
Neural variability does not encode interindividual sensory sensitivity but reflects intraindividual sensory perception. **(A)** Point-by-point partial correlations between trial-averaged SD ([high+ low)/2) and sensitivity measured by mean ratings ([high+ low]/2) to sensory stimuli of four modalities in Dataset l (N = 14l). No significant correlations were found in any modalities. Part-corr= Partial correlation. The light-colored “bursty.. curves represent the subject-averaged residuals of mean SD after regressing out mean amplitude. **(B)** Point-by-point intraindividual correlations between neural variability and ratings of sensory stimuli for each trial in Dataset I (N = 141). The gray bars represent *t* statistics at time points where significant differences were observed after raise discovery rate correction. **(C)** Point-by-point partial correlations between trial-averaged SD and mean ratings of sensory stimuli of four modalities in Datasets 2&3 (N =225). **(D)** Point-by-point intraindividual correlations between neural variability and ratings of sensory stimuli for each trial in Datasets 2&3 (N = 225). These results illustrate that neural variability encodes sensory perception at the intraindividual level but does not encode sensory sensitivity at the interindividual level.

## Discussion

Moving beyond the traditional amplitude-based approach, we systematically investigated the relationship between neural variability and pain, particularly pain discriminability, in five large EEG datasets (total N = 489). Five major findings were obtained. First, there were robust and replicable correlations between neural variability and pain discriminability across individuals in multiple datasets. Second, almost no significant correlations were observed between neural variability and sensory discriminability in tactile, auditory, and visual modalities, even when the perceptual ratings of nonpain modalities were matched with those of pain. Third, neural variability and ERP amplitude were mutually independent in encoding pain discriminability, and they had distinct temporal dynamics, efficiency, and underlying oscillatory profiles. Fourth, the association between neural variability and pain discriminability was partly disrupted on the pain-affected side of patients with PHN. Fifth, neural variability correlated with perceptual ratings in pain and nonpain modalities at the intraindividual level, but did not significantly correlate with individual differences in sensory sensitivity. Taken together, these results demonstrate that neural variability reliably and selectively encodes pain discriminability above and beyond amplitude, thereby enhancing the understanding of neural encoding of pain discriminability and facilitating a more comprehensive analysis of neural activity that incorporates both neural variability and ERP amplitude.

ERP amplitude is a central metric in most EEG studies [5,15,20,39,40]. However, neural activity is highly variable across time and such variability cannot be captured by the averaged amplitude itself [6,8,41]. Importantly, recent studies have linked neural variability to multiple cognitive processes, including perception and sensory discrimination [9,42,43]. Inspired by these discoveries, we moved beyond the traditional focus on the amplitude of neural activity and investigated how temporal neural variability of EEG responses relates to pain, in particular to interindividual pain discriminability. Consistent with recent studies showing an indirect connection between neural variability and pain [29,30], in our study neural variability reliably correlated with pain discriminability in multiple large datasets. This association remained robust across different measures of pain discriminability (i.e., rating differences and AUC) and different window sizes (i.e., 50 ms, 100 ms, and 200 ms). Interestingly, we also observed a selective correlation between neural variability and pain discriminability, as there were no reliable correlations between neural variability and tactile, auditory, and visual discriminability. Importantly, after matching intensity ratings between pain and nonpain modalities, we still found that neural variability significantly correlated with pain discriminability, but not with nonpain discriminability, thus effectively ruling out the potential confounding of intensity-rating differences between modalities. This replicable and selective association between neural variability and pain discriminability parallels the similarly robust and selective relationship observed between ERP amplitude and pain discriminability [19,20]. Moreover, similar to ERP amplitude, neural variability did not significantly correlate with pain sensitivity across individuals, but it did correlate with intraindividual perceptual ratings across trials [15]. These findings highlight that neural variability and ERP amplitude could have similar roles in encoding pain, and suggest that neural variability and ERP amplitude should both be considered in pain studies.

SD and mean values are associated mathematically. One could thus argue that the observed correlations between neural variability (measured by temporal SD) and pain may be simply a manifestation of the underlying relationships between ERP amplitude (measured by temporal means) and pain [15,20]. However, three lines of evidence speak against this notion. First, we observed partial correlations between neural variability and pain discriminability after controlling for ERP amplitude. Second, we found significant correlations between pain discriminability and another information-theory-based measure of neural variability, namely, PE, which is less influenced by the absolute values of ERP amplitude [35]. Third, the correlations between neural variability and pain discriminability remained in induced EEG responses, which refer to the component of neural responses independent of evoked activity (i.e., ERPs) [36]. Therefore, the correlations between neural variability and pain discriminability are unlikely to be solely amplitude driven but rather, attributable to neural variability itself. These findings agree with previous research demonstrating the independent contributions of variability and amplitude to cognitive processes [44–46].

Furthermore, we found differential roles and oscillatory profiles for neural variability and ERP amplitude as indicators of pain discriminability. Neural variability accounted for pain discriminability as well as or even better than ERP amplitude in the early time window following stimulus onset. Conversely, ERP amplitude seemed more crucial in the later window. These observations may be explained by SD’s sensitivity to small fluctuations in neural activity compared to means. Furthermore, fewer trials were needed to establish significant correlations between neural variability and pain discriminability. Neural variability thus seems more efficient in encoding pain discriminability, possibly because it can be measured relatively accurately at the single-trial level. Crucially, distinct oscillatory profiles were found for the roles of neural variability and ERP amplitude in encoding pain discriminability. While ERP amplitude was mainly observed in delta and theta frequencies, neural variability involved activity in broader frequency ranges: not only delta and theta bands, but also alpha and beta bands where ERP responses were absent. This result agrees with those of previous studies showing that neural variability may be a consequence of oscillations in alpha-beta bands [47]. Taken together, these findings provide further evidence that the relationship between neural variability and pain discriminability cannot be attributed to ERP amplitude. Instead, neural variability constitutes an independent and critical aspect of EEG responses with distinct mechanisms.

The independent and different roles of neural variability and ERP amplitude underscore the notion that the “noise” of neural activity is not purely noise but rather functionally informative [48]. Single-trial EEG signals typically appear very “noisy,” varying dramatically across time and even suppressing ERP responses [49,50]. This temporal variability is treated as noise and averaged out in the traditional trial-averaging procedure [51]. However, our findings show that this fluctuating temporal dynamics contains information about pain discriminability. This point is particularly illustrated by the “bursty” residual time series of neural variability after regressing out amplitude: the residual neural variability differed in the high- and low-pain conditions and correlated with pain discriminability. Our findings are in accord with those of previous studies associating task-related neural variability with sensory perception. For example, neural variability has been correlated with the visual contrast discrimination threshold [42], and the variability of blood-oxygen-level-dependent signals has shown correlations with discriminability in touch and visual motion [9,42,43]. Notably, most previous task-based studies examined trial-by-trial neural variability, that is, how neural activity varies across trials [42,47,52]. We, on the other hand, focused on temporal neural variability—moment-to-moment fluctuations in neural activity—which has largely been examined in the context of resting-state research [6]. Our study thus extends this line of research by demonstrating that moment-to-moment neural variability responds to external stimuli and is related to pain within task-based paradigms.

Why does temporal neural variability selectively encode pain discriminability? One possible explanation is that pain is a warning signal, indicating potential or actual harm to the physiological state. Therefore, pain perception may exhibit a higher level of alertness and attention than nonpain perception [53,54], which could be manifested by more pronounced variations in neural activity [41]. Indeed, neural variability can be generated by changes in attention and arousal levels, reflecting the richness of stimuli and difficulty of the cognitive process [41,55,56]. Two observations in our study seem to support this explanation. First, the neural variability differences between high- and low-intensity painful stimuli (ΔSD) seemed greater than or equal to those between high- and low-intensity nonpainful stimuli, despite the smaller rating differences for painful stimuli compared to nonpainful stimuli. Second, after the intensity-rating-matching procedure, neural variability for pain had a wider range than that for nonpain. In other words, pain seems to evoke greater neural variability. Another potential explanation could be the inherent unpleasantness of painful stimuli. In contrast to tactile, auditory, or visual stimuli, painful stimuli are inherently unpleasant [57–60]. Consistent with this notion, selective processing of pain has been found in the medial dorsal nucleus of the thalamus and its functional connectivity with the dorsal anterior cingulate cortex (ACC) and insula [61], which are implicated in the affective-motivational aspect of pain [58]. In support of this explanation, the topography of correlations between neural variability and pain discriminability showed a largely central distribution, which is in accord with neural sources in the ACC and insula [62]. Future studies can directly assess this hypothesis by examining whether neural variability also reflects discrimination of emotional stimuli, such as aversive sounds or itch stimuli [63,64]. Note that these two explanations are not mutually exclusive. Indeed, neural variability exhibited correlations with pain discriminability in a wide time range, which could involve both early-stage processing such as alertness and late-stage processing such as emotion.

In addition to the theoretical value of introducing temporal neural variability as an independent indicator of pain discriminability, the generalizability and potential clinical implications of our study are noteworthy. In Dataset 4 with multiple levels of stimuli, we observed stable correlations between neural variability and pain discriminability when stimulus intensity differences were moderate (i.e., 0.5 J), but the correlations became unstable when stimulus intensity differences increased. Similar findings were obtained in patients with PHN in Dataset 5, but the stimulus intensity differences were slightly greater (i.e., 0.75 J). The difference could be an aging effect, as the patients were older than the healthy participants in Dataset 4, and elderly persons generally have poor pain discriminability [65,66]. Crucially, the associations between neural variability and pain discriminability were disrupted on the chronic-pain-affected side, suggesting potential clinical relevance of neural variability as an indicator of pain discriminability. Previous studies have also revealed that patients with chronic pain exhibit abnormal pain discriminability, which correlates with the effectiveness of their treatment [23–26,67,68]. Validating our findings in different types of chronic pain will offer further implications for the diagnosis and prognosis of chronic pain, as well as for developing optimized treatment strategies based on neural variability as a pain discriminability indicator.

Despite its theoretical importance and potential clinical relevance, our study has some limitations. First, although replicable and selective correlations were found between neural variability and pain discriminability, the causality of this relationship remains unclear. It is likely that neural variability merely mirrors rather than determines pain discriminability. Further investigations employing other techniques, such as neurophysiology and neuromodulation, are needed to examine the causal direction. Second, the current study used only one type of pain, namely, laser heat pain. Previous studies have suggested different neural mechanisms across different types of pain [69]. It is thus essential to explore whether neural variability encodes the discriminability of alternative forms of pain, such as mechanical pain or chemical pain. Third, the clinical implications of our findings need more testing in patient groups. While we did test the ability of neural variability to serve as an indicator of pain discriminability in Dataset 5, it is only a small clinical dataset with only one type of chronic pain. Without broader testing in patients, the clinical relevance of neural variability is still limited. Addressing these limitations, future studies will deepen the understanding of how neural variability relates to pain and facilitate valuable clinical applications of neural variability.

## Methods

### Datasets and participants

Five EEG datasets were utilized in this study [15,20,70]. Datasets 1–4 comprised signals recorded from 462 healthy participants who were pain-free and had no history of chronic pain, neurological disorders, or psychiatric disorders. Dataset 5 was collected from 29 patients with PHN (2 were excluded in this study owing to insufficient trials for calculating the neural variability difference). Dataset 1 was recorded using the BioSemi EEG system, while Datasets 2–5 were recorded using the Brain Products EEG system. The detailed demographics are as follow: (1) Dataset 1 was collected from 141 participants (54 males) aged 21.8 ± 4.7 years (mean ± SD); (2) Dataset 2 was collected from 111 participants (54 males) aged 20.9 ± 2.3 years; (3) Dataset 3 was collected from 114 participants (40 males) aged 20.7 ± 2.3 years; (4) Dataset 4 was collected from 96 participants (51 males) aged 21.6 ± 1.7 years; and (5) Dataset 5 was collected from 27 participants (12 males) aged 61.7 ± 11.1 years. All participants gave written informed consent and were paid for their participation. The experimental procedures for Datasets 1–3 were approved by the local ethics committee at the Institute of Psychology, Chinese Academy of Sciences, and the procedures for Dataset 4 were approved by the local ethics committee at Southwest University (China). Dataset 5 was approved by the local ethics committee at Affiliated Daping Hospital of the Third Military Medical University (China).

### Experiments

Experimental procedures were identical for Datasets 1–3 (shown in Figure 1A). Participants were seated in a comfortable position in a dimly lit, quiet, and temperature-regulated environment. The entire experiment had three blocks, with each block comprising the administration of 40 sensory stimuli (encompassing 5 stimuli for every modality and intensity) while simultaneously recording EEG data. Participants were granted short breaks between blocks. Notably, the sequence of stimulus modality and intensity within each block was pseudorandomized. Each trial started with a fixation cross displayed for 3 s, followed by a transient sensory stimulus. After a 3-s interval, participants were asked to verbally rate the perceived intensity on a 0–10 numeric rating scale (NRS) within 5 s. A subsequent trial would initiate after 1–3 s following the rating period, resulting in an interstimulus interval ranging from 12 to 14 s.

For Dataset 4, participants received 40 stimuli, each with one of four stimulus intensities (i.e., 2.5 J, 3.0 J, 3.5 J, and 4.0 J), consisting of ten laser pulses per intensity. The sequence of stimulus intensity was pseudorandomized as well. The interstimulus interval varied randomly within 10–15 s. An auditory tone was presented 3–6 s after the stimulus, prompting participants to rate the intensity of the pain sensation elicited by the laser stimulus using the 0–10 NRS.

A similar experiment with multiple levels of stimulus intensity was performed on patients for Dataset 5. For each patient, ten laser pulses at one of six stimulus intensities (i.e., 3.0 J, 3.25 J, 3.5 J, 3.75 J, 4.0 J, and 4.25 J) were respectively delivered on each side (i.e., pain-affected or unaffected side). The order of stimulus energies was pseudorandomized, and the interstimulus interval varied randomly between 10 and 15 s. The participants rated the intensity of pain sensation following each laser stimulus using the 0–10 NRS.

### Sensory stimulation

For Datasets 1–3, participants received transient stimuli of four sensory modalities: nociceptive laser, non-nociceptive tactile, auditory, and visual. Each sensory modality encompassed two levels of stimulus intensity (i.e., high and low).

Following each stimulus, participants were asked to verbally rate the perceived intensity on an NRS from 0 (“no sensation”) to 10 (“the strongest sensation imaginable”). The intensities of tactile, auditory, and visual stimuli were carefully set based on a pilot experiment, aiming to elicit perceived ratings of approximately 4 and 7 out of 10 for low and high intensities, respectively. However, because of the potentially unbearable nature of the 4.0 J nociceptive laser stimulus for certain participants, a categorizing approach was adopted. Specifically, participants were categorized into high and low pain sensitivity groups. The high pain sensitivity group comprised individuals who rated the 4.0 J laser stimulus as above 8 out of 10 in the pilot experiment (Dataset 3). Conversely, participants who rated the 4.0 J stimulus as 8 or below were assigned to the low pain sensitivity group (Datasets 1 and 2).

The painful stimuli were delivered as short pulses of radiant heat (wavelength: 1.34 μm; pulse duration: 4 ms), generated by an infrared neodymium yttrium aluminum perovskite (Nd: YAP) laser (Electronical Engineering, Italy). The laser beam, transmitted through a 7-mm-diameter optic fiber, targeted a predefined 5 × 5-cm² area. To mitigate the risk of nociceptor fatigue or sensitization, the placement of the laser beam was randomly adjusted by about 1 cm after each pulse. For Datasets 1 and 2, which were collected from participants with low pain sensitivity, stimulus energies of 3.5 J and 4.0 J were used. For Dataset 3, involving participants with high pain sensitivity, the stimulus was delivered with stimulus energies of 3.0 J and 3.5 J. For Datasets 4 and 5, participants received nociceptive laser stimuli at four and six intensity levels, respectively.

Non-nociceptive tactile stimuli consisted of constant-current square-wave electrical pulses (with a duration of 1 ms from model DS7A, Digitimer, UK), delivered through skin electrodes positioned 1 cm apart on the left wrist, over the superficial radial nerve. The same two stimulus intensities were used for all three datasets: 2.0 mA and 4.0 mA. Auditory stimuli consisted of brief pure tones (with 800-Hz central frequency, 50-ms duration, and 5-ms rise and fall times) delivered through headphones. All participants were exposed to the same two intensities (76- and 88-dB sound pressure level). Visual stimuli took the form of brief flashes of a gray round disk presented on a black background (with a duration of 100 ms) on a computer screen. The intensities were adjusted using the gray scale of the disk, with RGB values set at (100, 100, 100) and (200, 200, 200).

### EEG recording and preprocessing

EEG data were acquired via 64 AgCl electrodes positioned according to the International 10–20 System, using the nose as the reference (band-pass filter: 0.01 to 100 Hz; sampling rate: 1000 Hz; BioSemi EEG system, Netherlands for Dataset 1; Brain Products EEG system, Germany for Datasets 2–5). Electrode impedance was kept below 10 kΩ. Electrooculographic signals were simultaneously recorded using two surface electrodes, one placed ∼10 mm below the left eye and the other placed ∼10 mm from the outer canthus of the left eye.

EEG data were preprocessed in MATLAB (R2021b; MathwWorks, USA) using the EEGLAB toolbox v2023.1 [71]. Continuous EEG data were band-pass filtered between 1 and 30 Hz. Subsequently, EEG data were segmented into epochs extending from 500 ms before to 1000 ms after stimulus onset. Each epoch was baseline corrected using the prestimulus interval. Trials contaminated by eye blinks and movements were corrected using an independent component analysis algorithm implemented in the EEGLAB toolbox [71].

### Neural variability

We mainly used temporal SD, a variance-based neural variability measure at the single-trial level and then averaged across trials of the same stimulus intensity. Temporal SD is defined as the standard deviation of EEG responses within a time window:

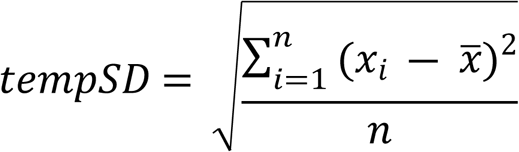

where *x*_*i*_ is the amplitude at time point *i*, and *x* is the mean amplitude within the time window of *n* time points. We applied the sliding window of 100 ms (*n* = 100) to delineate how temporal SD changes over time (as shown in Figure 1B). Additionally, to evaluate the impact of time window duration on the results, we repeated analyses using time windows of 50 ms and 200 ms.

We also used the information-theory-based measure PE as an additional metric of neural variability to demonstrate the robustness of the relationship between neural variability and pain discriminability. PE quantifies the complexity of a time series by evaluating the order relations between values in the time series rather than their absolute numerical values [35]. We computed PE with the customized function *PE* from the MATLAB file exchange [72]. The computation process involves selecting an embedding dimension *m*, representing the number of sequential time points forming a pattern, and a time delay *τ*. Then vectors 𝑣_*t*_ = (*x*_*t*_, *x*_*t*+*τ*_, *x*_*t*+2*τ*_, …, *x*_*t*+(*m*−1)*τ*_) are extracted from a windowed time series (*x*_1_, *x*_2_, …, *x*_*n*_), where *n* denotes the number of time points within the specified window. These vectors facilitate the enumeration of relative frequencies for all potential ordinal patterns by ranking the elements to construct a probability distribution *P* across all patterns. Then PE can be calculated using Shannon entropy as follows:

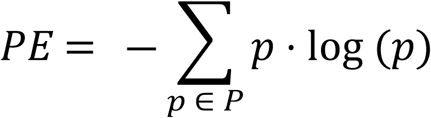

In our case, we set *m* = 4, *τ* = 1, and *n* = 100. Therefore, 97 vectors can be extracted from a time window of 100 time points, namely, 𝑣_1_ = (*x*_1_, *x*_2_, *x*_3_, *x*_4_), 𝑣_2_ = (*x*_2_, *x*_3_, *x*_4_, *x*_5_), …, 𝑣_97_ = (*x*_97_, *x*_98_, *x*_99_, *x*_100_). We then determined the relative frequencies of all possible ordinal patterns within these vectors, such as (1, 2, 3, 4), (1, 2, 4, 3), (1, 3, 2, 4), and so on and calculated Shannon entropy to obtain PE for each time window. As we did for the temporal SD calculation, we also employed the sliding window technique to characterize the temporal changes in PE.

### Sensory discriminability

Sensory discriminability was quantified by the difference between ratings averaged across trials of high intensity and trials of low intensity. A previous study has shown that this simple index has a less skewed distribution and yields results highly similar to those of other indices, such as measures based on signal detection theory [20]. To underscore the robustness of our findings, we also conducted an assessment utilizing the index derived from signal detection theory in this study.

### Correlation analysis

To determine if neural variability reflects pain discriminability, we computed neural variability (i.e., temporal SD) at the single-trial level, took the difference of neural variability between two intensities (high–low, ΔSD), and correlated this neural variability difference with pain discriminability in a point-by-point manner in Datasets 1–3. Multiple comparisons were corrected using the FDR procedure [73]. Scalp topographies were computed by spline interpolation. To illustrate the correlations in scatterplots, we extracted the values within a 20-ms window centered at the peak of the subject-averaged neural variability difference and correlated them with pain discriminability.

We also correlated mean neural variability ([high + low]/2) with mean pain ratings to determine if neural variability also reflects interindividual pain sensitivity. Neural variabilities were computed at the single-trial level and averaged across all trials for each subject, regardless of stimulus intensities. The pain sensitivity was quantified using the ratings averaged across all trials for each subject.

To explore the intraindividual trial-by-trial relationship between brain responses and pain intensity ratings, we computed correlations between single-trial neural variability and the corresponding ratings of pain intensity for each subject. The obtained correlation coefficients were transformed to *z* values using the Fisher *z* transformation, and the *z* values were finally compared against zero using a one-sample *t* test. All statistical tests were two-tailed.

### Independence between variance-based neural variability and amplitude of brain responses

Since variance and amplitude of the brain response are generally not independent, we first correlated neural variability differences and amplitude differences (also referred to as ΔSD and ΔAmplitude, respectively) between high- and low-intensity conditions and then performed a partial correlation analysis between ΔSD and pain discriminability, controlling for ΔAmplitude at the same time point. Note that, consistent with neural variability, ERP amplitude was also calculated with a 100-ms sliding window. To further account for ΔAmplitude, we subtracted the trial-averaged amplitude from single-trial EEG responses for each condition to derive induced EEG responses [36]. We then computed neural variability and conducted correlation analysis with the induced EEG responses. To illustrate the correlations in scatterplots, we also extracted the values within a 20-ms window centered at the peak of the subject-averaged neural variability difference. After regressing out the ΔAmplitude, we created scatterplots using the residuals of ΔSD and pain discriminability.

### Dominance analysis

To assess the contributions of differential neural variability and differential amplitude (i.e., ΔSD and ΔAmplitude) to encoding pain discriminability, we conducted a dominance analysis at each time point [37,38]. Dominance analysis is a method for comparing the relative contribution of predictors in multiple regression models. We quantified the contribution using total dominance, which has the desirable property that the sum of total dominance of all predictors is equal to the determination coefficient (*R*^2^) of the full multiple regression model. At each time point, we first regressed pain discriminability against differential neural variability and differential amplitude and computed the *R*^2^ of this full model, which is denoted 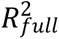. Then, we predicted pain discriminability separately using differential neural variability and differential amplitude. The *R*^2^s of these two reduced models are denoted 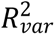 and 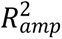. According to the dominance analysis approach, total dominance of neural variability differences is defined as 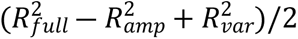. Similarly, total dominance of amplitude differences is 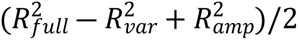.

### Resampling approach

We used a resampling method to examine the influence of number of subjects and number of trials on the probability of detecting significant correlations between neural variability and pain discriminability. In Datasets 1–3, we generated bootstrapped subsamples from the whole dataset, that is, randomly selected data samples with replacement. For the number of subjects, we subsampled subjects ranging from 20 to the maximum of each dataset and repeated the resampling procedure 100 times for each number of subjects. To assess the effect of number of trials, we subsampled trials ranging from 1 to 15 (in steps of 1), and for each number of trials, the resampling procedure was repeated 100 times. Note that during the resampling of subjects, we used data from all trials, while during the resampling of trials, data from all participants were used. In each subsample, we computed differential neural variability and differential ERP amplitude, correlated them with pain discriminability, and applied a threshold to the correlation time series with *p*(FDR) = 0.05. Since we generated 100 subsamples for each sample size, the probability of a time point being significant under a certain number of subjects/trials was approximated by the number of times that time point reached significance. We also specifically examined the number of subjects/trials needed to detect significant correlations with a probability ≥ 80% [74].

### Oscillatory profiles of neural variability and ERP amplitude

To further explore whether neural variability and ERP amplitude were driven by frequency-specified activity, we band-pass filtered single-trial EEG signals to characterize the oscillatory profiles of neural variability and ERP amplitude as indicators of pain discriminability. Since continuous EEG traces were filtered from 1 Hz to 30 Hz to derive the ERPs, here we focused on four classic frequency bands: delta (1–4 Hz), theta (4–8 Hz), alpha (8–12 Hz), and beta (12–30 Hz). After band-pass filtering the data for the respective frequency band, we then repeated the partial correlational analyses, that is, correlating ΔSD in the filtered data with pain discriminability in a point-by-point manner while controlling for ΔAmplitude at the same time point, and correlating ΔAmplitude in the filtered data with pain discriminability while controlling for ΔSD at the same time point.

### Pain selectivity of neural variability for encoding sensory discriminability

To test the pain selectivity of the relationship between neural variability and sensory discriminability, we repeated correlation analyses and partial correlations controlling for ERP amplitude on Dataset 1–3 with sensory stimuli of tactile, auditory, and visual modalities.

Perceptual ratings can vary significantly between different sensory modalities, making it challenging to compare the relationship between EEG responses and discriminability indices across modalities. This variability in perceptual ratings might confound any observed differences between modalities. To account for this potential confounder, we used a rating-matching procedure to equalize intensity ratings between pairs of sensory modalities (i.e., pain vs. touch, pain vs. audition, and pain vs. vision) in both Dataset 1 and Dataset 2&3.

The matching procedure was adapted from a previous study [75]. When matching pain and touch ratings, if Participant X rates high- and low-intensity laser stimuli as, for instance, 6 and 4 on average, respectively, we identify participants whose mean ratings for high-intensity tactile stimuli fall within 5.5 to 6.5 and for low-intensity tactile stimuli within 3.5 to 4.5. This ensures an absolute average matching error of ≤ 0.5 for both high- and low-intensity stimuli. If M participants meet this criterion, we further narrow it down to N participants with the smallest absolute matching error. If N equals 1, Participant X is paired with this single participant. If N is greater than 1, Participant X is randomly paired with one of these participants (say, Participant Y), provided that Participant Y has not been paired with anyone else. The effect of this matching procedure was confirmed by statistically insignificant interaction effects (sensory modality × stimulus intensity) in a mixed-design ANOVA.

### Pain discriminability of multiple intensities in healthy subjects and chronic pain patients

We explored the correlations between neural variability differences and pain discriminability in Datasets 4 and 5, both recorded from participants exposed to various intensities of nociceptive stimuli. In Dataset 4, four levels of stimuli with a 0.5-J interval were used, resulting in six pairs of high- and low-intensity combinations. We first applied a one-way repeated-measures ANOVA to determine whether ratings of pain intensity differed across stimulus intensities. We then performed paired-sample *t* tests on temporal SDs evoked by various pairs of stimulus intensities. For each pair of stimulus intensities, we performed a correlation analysis as previously described between neural variability and pain discriminability. In Dataset 5, six levels of stimuli separated by a 0.25-J interval were applied on both the affected and unaffected sides, resulting in 15 pairs of high- and low-intensity combinations for each side. We first applied a two-way repeated-measures ANOVA to determine whether ratings of pain intensity were significantly different across stimulus intensities and stimulation sides. We then performed paired-sample *t* tests on ratings and temporal SD values of different pairs of stimulus intensities. A correlation analysis was then conducted on those pairs of high- and low-intensity combinations that showed significant differences in both ratings and temporal SDs.

## Data and code availability

The data and code for all results will be made available on the Open Science Framework (OSF) upon publication.

## Acknowledgement

This work was supported by the National Key Research and Development Program of China (2023YFC2508702), National Natural Science Foundation of China (32071061), and Beijing Natural Science Foundation (JQ22018; L246074).

## Author Contributions

Conceptualization: LBZ, LH, XYG; Methodology: LBZ, LH, XYG; Investigation: LBZ; Formal analysis: XYG, LBZ; Visualization: XYG, LBZ, LH; Writing – original draft: LBZ, XYG; Writing – review & editing: LBZ, LH, XYG; Funding acquisition: LH; Project administration: LBZ, LH; Supervision: LH.

